# Investigating the potential of opportunistic sighting data to inform wildlife conservation strategies

**DOI:** 10.1101/075945

**Authors:** Simone Tenan, Paolo Pedrini, Natalia Bragalanti, Claudio Groff, Chris Sutherland

## Abstract

Abundance and space use are key population-level parameters used to inform management and conservation decisions of rare and elusive species, for which monitoring resources can be limited, potentially affecting quality of model-based inference. Recently-developed methods that integrate multiple data sources arising from the same ecological process have typically been focused on data from well-defined sampling protocols, i.e. structured data sets. Despite a rapid increase in availability of large datasets, the value of unstructured or opportunistic data to improve inference about spatial ecological processes is, however, unclear. Using spatial capture-recapture (SCR) methods, we jointly analyze opportunistic recovery of biological samples, traditional SCR data resulting from systematic sampling of hair traps and rub trees, and satellite telemetry data, collected on a reintroduced brown bear population in the central Alps. We compared the precision of sex-specific estimates of density and space use derived from models using combinations of data sources ranging from traditional SCR to a fully integrated SCR model that includes both telemetry and opportunistic data. Estimates of density and space use were more precise when unstructured data were added compared to estimates from a classical SCR model. Our results demonstrate that citizen science data lend itself naturally to integration with in the SCR framework and highlight the value of opportunistic data for improving inference about space use, and in turn, of abundance and density. When individual identity and location can be obtained from opportunistic observations, such data are informative about space use and thus have the potential to improve estimates of movement and density using SCR methods. This is particularly relevant in studies of rare or elusive species, where the amount of SCR encounters is usually small, but also budget restrictions and the difficulty of collaring animals limit the number of individuals for which telemetry information is available. Spatially-referenced opportunistic data thus potentially increase both the geographic extent of a study and the number of individuals with available spatial information, providing an improved understanding of how individuals are distributed and how they use space – fundamental components for calibrating conservation management actions.

## INTRODUCTION

Obtaining precise estimates of population density and space use can lead to an increased understanding of the processes governing spatio-temporal ecological dynamics and, in turn, improve wildlife management and conservation practices. The task of estimating ecological state variables is, however, challenging, especially for rare and elusive species such as large carnivores, and requires analytical approaches that account for the fact not all individuals in a population can be observed (Williams et al. 2002). Regardless of methodology, the quality of model-based inference is directly related to data quality, which can be an issue for elusive species, especially when resources for monitoring are limited. This has led to an emphasis on developing methods that integrate multiple data sources (Schaub and Abadi 2010; Gimenez et al. 2014) and, importantly, to a realization that the vast amounts of data regularly collected outside of formal scientific studies, *unstructured* or *opportunistic* data, offer a potentially valuable data source (Dickinson et al. 2012; Newman et al. 2012). Although the majority of data integration methods have focussed on improving estimates of species distribution and temporal population trends, opportunistic data has great potential to improve inferences about spatial ecological processes.

Integrated population models (IPMs: Besbeas et al. 2002; Schaub and Abadi 2010; Tenan et al. 2012) provide a statistical framework for jointly modeling count data and demographic data, typically resulting in improved inferences about the mechanisms regulating population dynamics. As a result, there has been continued development of more general ‘integrated data models’ that seek to combine any independent data sources that arise from the same ecological process (Gimenez et al. 2014). For example, occupancy and abundance are two directly related ecological state variables and joint analysis of capture-recapture and occupancy data has been shown to improve estimates of abundance (Conroy et al. 2008; Blanc et al. 2014), density (Chandler and Clark 2014), and even colonization-extinction dynamics and dispersal (Sutherland et al. 2014). A common feature of the majority of studies that use multiple data sources, aside from improving parameter precision, is that each independent data set is collected according to a well-defined sampling protocol, i.e., it is an integration of structured data sets. The value of unstructured or opportunistic data, such as that collected by many citizen scientists, is yet unclear. For example, van Strien et al. (2013) argue that opportunistic data represents an important data source that, if analyzed appropriately, can yield improved inferences about temporal trends in occurrence, while Kamp et al. (2016) caution against its use, demonstrating that citizen science data were unable to detect significant species declines. Regardless, with the rapid increase in citizen science initiatives, finding innovative ways to utilize opportunistic data will broaden the scope of ecological enquiry that can be addressed within a single analytical framework (Gimenez et al. 2014).

Spatial capture-recapture methods (SCR: Efford 2004; Royle et al. 2014) are now well-established in applied ecology and produce estimates of population density using spatial encounter history data, i.e., observation data on who was detected when, and importantly, where. Using spatial patterns of observations to account for heterogeneity in detection probabilities caused by individual differences in trap exposure, and treating space as an explicit model component, SCR produces unbiased estimates of density and space use across a range of conditions (e.g., Borchers and Efford 2008; Royle et al. 2014; Sollmann et al. 2012; Sutherland et al. 2015). Moreover, SCR has been used to estimate density for elusive species from data collected using a variety of field methodologies including camera traps (Royle et al. 2009), hair snares (Gardner et al. 2010), scat surveys (Fuller et al. 2016), and unstructured survey data (Kéry et al. 2011). A core component of SCR is an explicit model for space use that relates encounter probability to the distance between any location and an individual’s activity center via the estimation of a spatial scale parameter *σ* (Ch. 7 Royle et al. 2014). Estimating *σ* accounts for individual encounter heterogeneity so the effective sampling area is explicitly described and as a result, absolute density can be directly estimated. If follows that to estimate density well, *σ* must also be well estimated. As with other statistical methods, the precision of SCR-derived estimates of space use and density depend on sample sizes, specifically, but not solely, the number of unique spatial locations individuals are observed at (spatial recaptures). Thus, adding additional spatial information should, in theory, lead to improved inference about space use, and in turn, density. For example, Gopalaswamy et al. (2012) increased the number of spatial recaptures by integrating camera trap and scat collection data which resulted in more precise estimates of density, while Royle et al. (2013) and Sollmann et al. (2013c) demonstrated that space use (*σ*) and density are estimated with higher precision when telemetry data are used in addition to traditional capture-recapture data (See also Table 1).

**Table 1:**
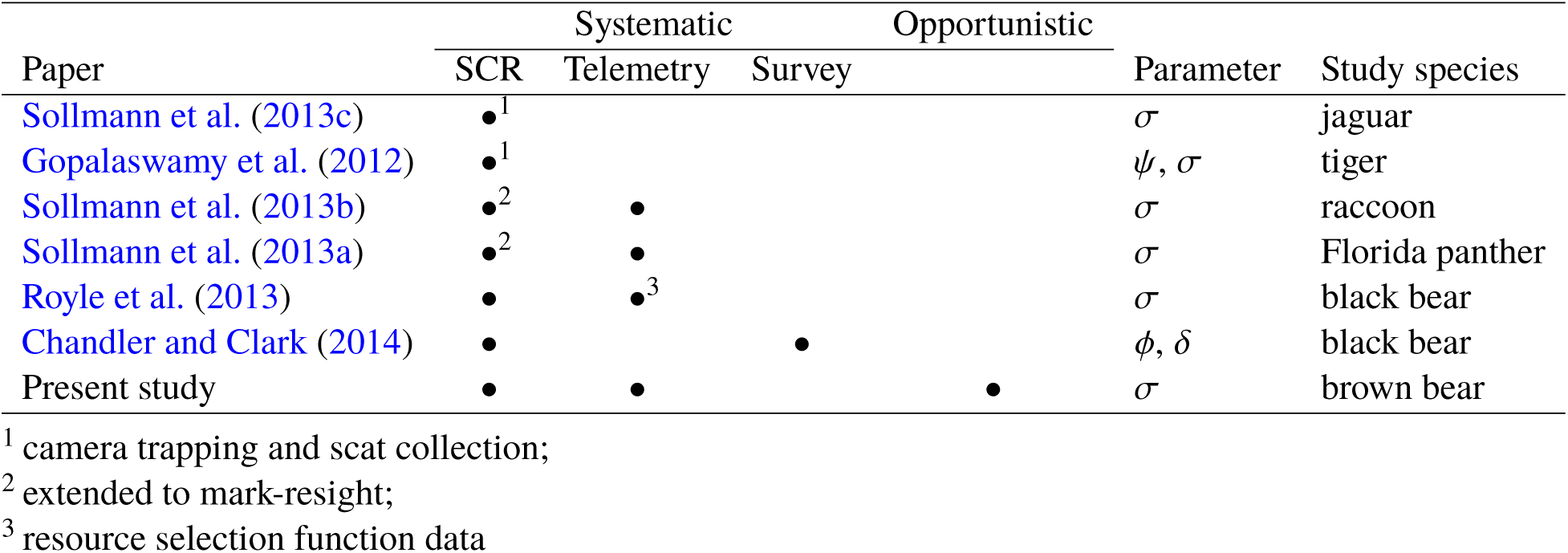
Summary of contributions that provide an integrated framework for spatially-referenced individual data. Systematic data are collected under specific study designs: spatial capture-recapture (SCR), telemetry, and counts or binary detections (survey). Parameter shared: *ψ*, Data Augmentation parameter; *σ*, scale parameter of the observation model; *φ*, survival probability; *δ*, individual-level recruitment probability.

Interestingly, in an SCR model, telemetry data require no information about sampling effort because observed locations provide representative information only about the spatial scale parameter (*σ*), and thus any amount of telemetry data are likely to be informative about space use. So, while the inability to quantify observer effort and bias is often cited as a major limitation of data collected by citizen scientists (Dickinson et al. 2010), it appears that such opportunistic data lends itself naturally to integration within the SCR modeling framework. Specifically, when opportunistic observations can be made of individuals, i.e., via direct or indirect recognition of naturally marked, collared or tagged individuals or the collection of DNA yielding biological samples such as hair of faeces, the locations of those observations are informative about space use and therefore have the potential to improve estimates of spatial parameters in SCR, and have the added benefit of potentially increase the geographic extent of monitoring studies significantly.

Here we demonstrate how opportunistic data can be jointly analyzed using spatial capture-recapture methods to improve estimates of density and space use for a reintroduced brown bear *Ursus arctos* population in the central Alps. The area is one of the most populated regions to be occupied by brown bears (Chapron et al. 2014; De Barba et al. 2010b) meaning bear-human interactions are highly probable and any perceived threat is considered a key factor in determining the success or failure of the reintroduction (Mustoni et al. 2003). It is important that any conservation management decisions are based on the best available information, and every effort must be made to improve estimates of bear density and space use.

## MATERIALS AND METHODS

### Study area and population

This study was conducted in 2013 in the Italian Alps, an area characterized by a mosaic of natural and human-modified habitats, with a landscape fragmented by urban areas and roads. Elevation ranges from 65 m to more than 3900 m a.s.l., with submontane, montane and subalpine vegetation covering areas below 2000 m, and human population density concentrated below 1000 m (Mustoni et al. 2003). Between 1999 and 2002, nine bears (three males and six females, 3-6 years old) were released in the area as part of a reintroduction project to establish a self-sustained population (Dupré et al. 2000; Mustoni et al. 2003). At the time, the original brown bear population consisted of at least three animals, which were assumed to have died without any genetic exchange with the translocated bears and their progeny (De Barba et al. 2010a).

### Brown bear data

#### Non-invasive genetic sampling

Bear hair samples were collected form 99 hair traps and 89 rub trees. Hair traps consisted of a strand of barbed wired wound around trees at *c.* 50 *cm* above ground level enclosing an area of *c.* 25 *m*^2^ and scent lure was placed in the center (Woods et al. 1999). They were set from 15 May to 31 July, checked on five occasions and the number of days between occasions ranged from 3 to 10 days (Fig. 1). Rub trees, barbed wire wrapped around trees, were monitored during the same time period and were checked twice, first after six days and than after four days (Fig. 1). All hairs were collected during each visit to hair traps and rub trees so that only newly deposited hairs were collected in subsequent visits. Because the hair trap and rub tree data were collected according to a specific protocol, we refer to this structured data as traditional SCR data, or simply ‘SCR data’. In addition to the structured data collection, we also collected opportunistic hair and feces data (De Barba et al. 2010a,b; Groff et al. 2014; Tenan et al. 2016). Following notification by third parties (typically members of the public), opportunistic sampling of hair and feces was carried out by agency personnel at sites where bear damage occurred, e.g. depredation on livestock, beehives and/or crops (Tenan et al. 2016). We refer to this data as ‘opportunistic data’.

**Figure 1:**
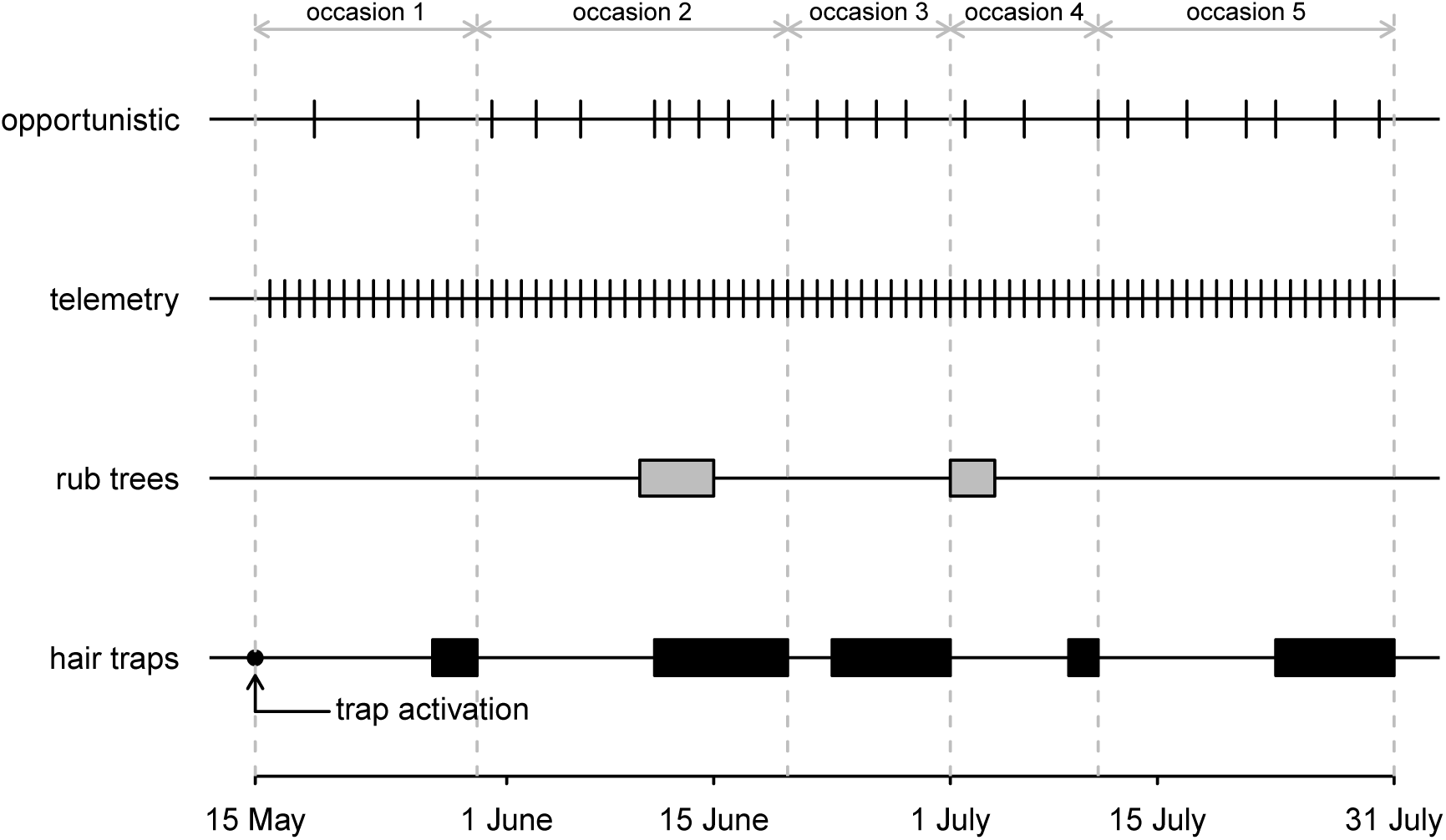
Timeline of data collection during the period when SCR data were systematically collected using an array of hair traps checked on five occasions of variable length (black blocks), and from rub trees checked for hairs in two period (grey blocks). Telemetry data were thinned by randomly selecting on record per day, and opportunistic recovery of biological samples was performed in 23 days.

Biological samples were genetically analyzed for individual identification using ten loci. For a detailed description of DNA extraction methods, PCR protocols, protocols for individual identification, and molecular sexing, see De Barba et al. (2010a,b). We considered only data belonging to the non-cub part of the population and successfully identified a total of *n* = 22 individuals (12 females and 10 males). Of the 22 individuals, 19 were detected using hair traps; two males and one female that were sampled only on rub trees. During the period of trap deployment, 11 of the 22 individuals (four females and seven males) were detected opportunistically resulting in an additional 30 unique spatial locations (Fig. 1, Fig. 2a, Fig. S1 in Appendix S1).

**Figure 2:**
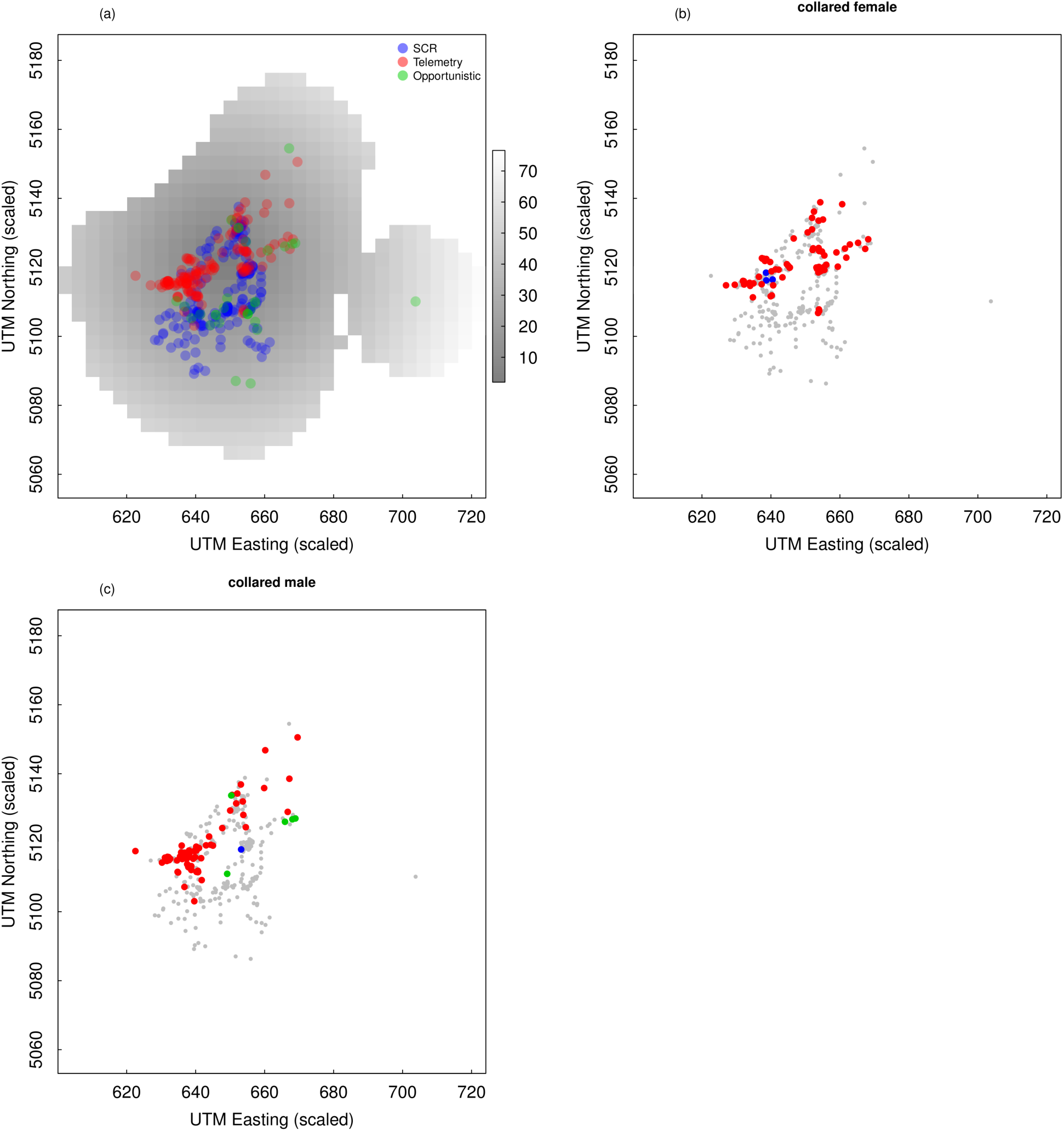
(a) Distance from the point were founders were released (in km) and location of bear captures from systematic sampling with hair traps and rub trees (SCR), telemetry and opportunistic records. (b-c) Location of the records for the two collared individuals from which telemetry information was derived. Grey dots indicate the location of all observed individuals.

#### Telemetry

Two bears, a 5-year old male and a 15-year-old female, were monitored during the hair trap and rub tree sampling period using Global Positioning System (GPS) collars (Vectronic GPS-GSM collars, Vectronic Aerospace GmbH, Berlin, Germany). Collared bears were captured using culvert traps (female) and Aldrich snares (male) upon approved capture protocols (2003-DPR 357/97, Groff et al. 2014). GPS collars collected positions at different intervals ranging from 10 min to 1 h. For the analysis we selected one random record per day per individual, giving a total of 143 unique telemetry locations (74 and 69 for the male and female respectively). The collared female was detected at a hair trap but never detected at rub trees or opportunistically (Fig. 2b). The collared male was never detected with hair traps, but was detected once at a rub tree and opportunistically observed in five occasions (Fig. 2c).

## Data analysis

### Spatial capture-recapture data

Spatial capture recapture models are hierarchical models (Royle and Dorazio 2008) that describe distance-dependent encounter probabilities (the observation process), and the spatial distribution of individuals across the landscape (density, the ecological state process). We adopt a Bayesian analysis of the model (Royle and Young 2008; Gardner et al. 2010) and assume that individual encounter data, *y*_*ijk*_, representing whether or not individual *i* was detected in trap *j* in occasion *k*, are Bernoulli random variables with success probability *p*_*ijk*_, i.e., the encounter probability:

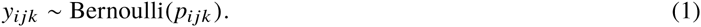

Encounter probabilities in SCR are assumed to decline with distance between a trap (*x*) and an individuals activity center (*s*) according to some decreasing function; here we use the commonly applied half-normal encounter model and allow for sex-specific variation in the parameters:

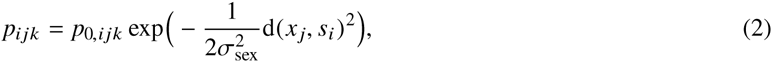

where *σ*_sex_ is the sex-specific spatial scale parameter that determines the decrease in encounter probability as the distance between trap *j* and individual *i*’s activity center (d(*x* _*j*_, *s*_*i*_)), increases. The parameter *p*0,_*ijk*_ is the baseline encounter probability and can itself be modelled as a function of individual-(*i*), trap-(*j*) and occasion-(*k*) specific covariates. Specifically, we modelled the baseline encounter probability as a function of *sex*, trap type (*trap*: hair trap or rub tree), and, to account for the different time elapsed between consecutive sample occasions in each trap, time since last check (*time*) using standard logistic regression:

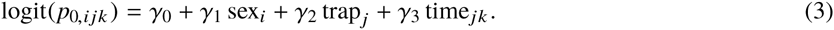

The second component of the SCR model is a point process model that describes the distribution of individual activity centers, *s*_*i*_, within a defined state-space S which should be large enough to contain all plausible activity centers of all observed individuals (Royle et al. 2014). We were particularly interested in modelling density as a function of spatially varying covariates, i.e., an inhomogeneous point process model, and so we used a discrete representation of S defined as the center points of each pixel. In our case, where we are considering a population that was established from a single release point, we modelled variation in density as a function of the distance from the point that the founding population was released between 1999 and 2002 (*d.release*; see ‘Study area and population’). Using a binomial point process model, the per-pixel intensity, *µ*(*s*), is modelled as a log-linear function of ‘d.release’:

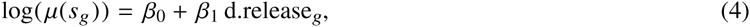

and the probability that an individual activity center is located in a pixel, *π*(*s*) is given by:

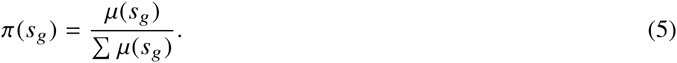

This is the standard formulation of a Bayesian SCR model with a sex-specific half-normal encounter probability model, an inhomogeneous point process density model, and the estimation of sex-specific total population size *N* using data augmentation (see Chapters 7 and 10 in Royle et al. 2014). Sex was known for all observed individuals but not for unobserved (augmented) individuals so was modelled as an individual random effect to be estimated: sex_*i*_ ∼ Bern(*ω*_*sex*_), where *ω*_*sex*_ is the population-level sex ratio. We expected detectability to vary significantly between sexes and trap type, and to be positively related to the time since last check, expected space use to vary by sex, and thus that sex-specific *σ* values would differ, and finally we expected density to decline with distance from the release point.

The spatial encounter histories for the standard SCR analysis were generated for detections from a *J* = 188-trap array consisting of hair traps and rub trees across *K* = 5 sampling occasions (Fig. 1 and Fig. 3). Data were formatted in a 3-dimensional *M* × *J* × *K* array, **Y**_*SCR*_, where *M* is the number of augmented ‘all-zero’ encounter histories, a proportion of which are the estimated unobserved individuals. The additional data required to fit the model are: the coordinates of each hair trap and tree rub, a vector of sex determination of each individual, a *J* × *K* trap operation matrix which is a binary indicator denoting whether each trap was operational during sampling occasion, and the *J* × *K* matrix of ‘time since last check’ covariates, which were scaled to have zero mean and unit variance (Fig. 3).

**Figure 3:**
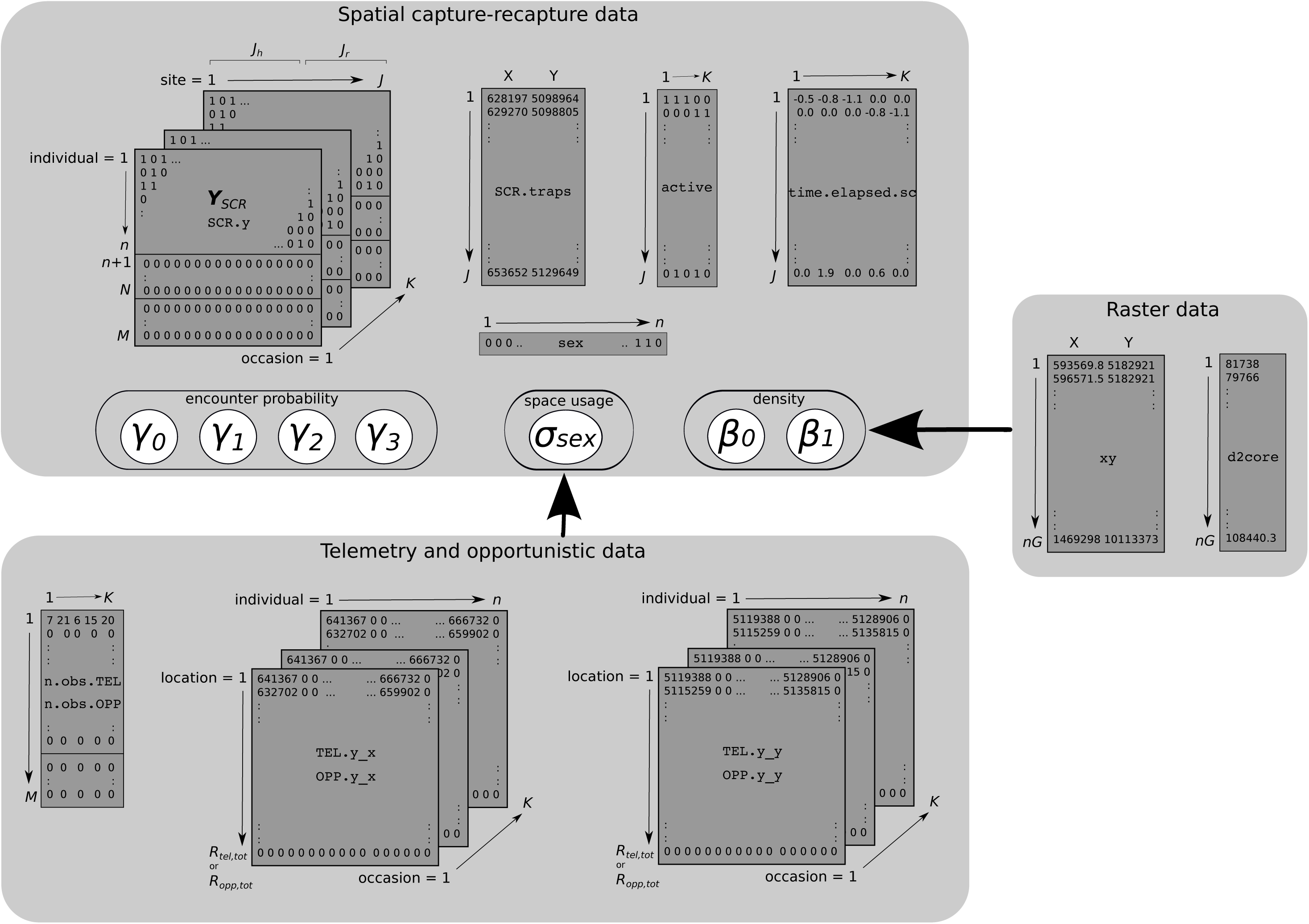
Graphical representation of the data involved in the integrated analysis. Circles represent estimated parameters. Observed data for *n* individuals, detected during *K* visits and members of the population of size *N*, were augmented with *M* − *n* all-zero detections (***Y*** _*SCR*_ matrix). SCR data were collected at *J* sites, consisting of *J*_*h*_ hair traps and *J*_*r*_ rub trees. Data set names in Courier font correspond to the names used in the model code. Coordinates, trap deployment, and (standardized) time since last check data sets for the *J* = *J*_*h*_ + *J*_*r*_ traps are denoted by SCR.traps, active, and time.elapsed.sc labels, respectively. Raster data contain information on the distance from the point were founders were released for each of the *nG* pixels (d2core) and was used to model density. Telemetry and opportunistic data were formatted in the same way, with an augmented matrix for number of records available for individual *i* in occasion *k* (n.obs.TEL and n.obs.OPP, respectively) and the x (TEL.y_x, OPP.y_x) and y (TEL.y_y, OPP.y_y) coordinates of those records for each individual in each of the *R*_*tel*_ or *R*_*opp*_ locations and occasion *k*.

### Telemetry and opportunistic data

Unlike the traditional SCR data described above, both telemetry locations, **I**_tel_ and opportunistic data **I**_opp_ are not restricted to trap locations and therefore provide important additional information about individual movement, i.e., they are both direct observations of space use (Royle et al. 2013). We combine the individual telemetry and opportunistic locations and refer to them collectively as **I**_*i*_ for individual *i* = 1*, …, n*. These additional locations can be modelled using a bivariate normal movement model (Sollmann et al. 2013b, a):

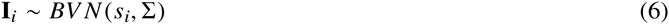

where,

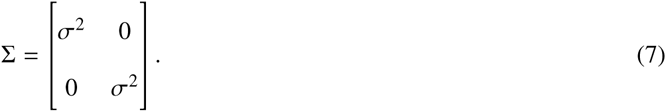

The parameters of this bivariate normal model can be related directly to the SCR half-normal encounter probability model (Eq. 2) through the shared parameters *s* an *σ*, which means that telemetry data, opportunistic data and traditional SCR data can be jointly modelled, each contributing to the estimation of the latent activity center, *s*_*i*_ location and the spatial scale parameter *σ*.

The telemetry and opportunistic data, for *R*_tel_ and *R*_opp_ locations, were formatted in two *R* × *n* × *K* arrays each, one containing x-coordinates and the other containing corresponding y-coordinates, for the *n* observed individuals and *K* occasions. This array structure allows the unstructured data (telemetry and opportunistic) to be related to the SCR data in the integrated model (Fig. 3). In addition, an *M* × *K* matrix denotes the number of unstructured locations for each individual in each occasion (Fig. 3).

### Model comparisons

The main objective of this work was to investigate the value of integrating telemetry data and opportunistic sightings data into a traditional SCR model for estimating density and space use for the reintroduced brown bear population to inform future management. To do so we fitted the SCR models described above using four SCR data sets:

1. Traditional SCR data only (data from hair traps and tree rubs)
2. SCR data + telemetry data
3. SCR data + opportunistic sightings
4. SCR data + telemetry + opportunistic sightings

For each model, we estimated sex-specifc total population size, *N*_sex_ and sex-specific *σ*_sex_, and compared point estimates (posterior median) and precision (95% Bayesian Credible Interval width, BCI from here) of the estimated parameters.

We adopted a Bayesian analysis of the SCR models using Markov chain Monte Carlo implemented using the program JAGS (Plummer 2003) executed from R (R Core Team 2012). For the parameters of the linear predictors (*γ* and *β*), we used an uninformative Normal(0, 100) prior. We used a equally uninformative Normal(0, 10) prior for *α*_1,sex_, where *α*_1,sex_ = 1/(2*σ*^2^_sex_). After testing a range of resolution values for the state space, we used a resolution of 4 × 4 *km*, a value that was small enough to yield stable parameter estimates, and large enough to ensure the model was computationally tractable. To ensure the state space was large enough to contain all plausible activity centers, we used a 21 *km* buffer around the most extreme coordinates of all the data (telemetry, opportunistic and trapping data, Fig. 2a). Data were augmented with *M* − *n* ‘all-zero’ encounter histories, where *M* = 300. Summaries of the posterior distribution were calculated from 30, 000 post-burn-in posterior samples (burn-in = 3, 000 iterations). The code for the fully-integrated model is available as supporting information.

## RESULTS

Overall, integrating all available sources of information (traditional SCR, telemetry and opportunistic data) produced more precise estimates of population size and spatial scale when compared to the use of either SCR data alone or integrated with only one additional source of data (Table 2–3, Fig. 4). In particular, the gain in precision achieved by jointly modeling all three data types was particularly relevant for sex-specific population size estimates (*N*_*m*_ and *N*_*f*_). In addition to the increase in precision, integrating additional sources of information resulted in a shift in the median abundance point estimates: from 15.679 (BCI: 7.994 – 30.349) under the SCR-only model to 12.629 (BCI: 6.840 – 21.953) under the fully integrated model for the number of males, and from 58.008 (BCI: 24.285 – 147.788) to 24.578 (BCI: 13.275 – 51.722) for females. Precision gains in estimates of *σ* were minimal when adding opportunistic data and highest when integrating telemetry information only, markedly so for males (Table 2–3, Fig. 4). As with the estimates of bear population size, the integration of additional information led to a change in the point estimates of *σ*; compared to the SCR-only model, there was a noticeable increase in the scale of space use when any of the additional data was used. The female 95% home range size estimated from the half normal encounter model under the SCR-only model was 212 *km*^2^ whereas for the fully integrated model, the estimate was 1375 *km*^2^. Conversely, estimates of male space use, *σ_m_*, were consistent across models (Fig. 4, Table 2), as were the corresponding 95% home range size estimates: 1800 *km*^2^ (SCR data only) and 2024 *km*^2^ (fully integrated model).

**Table 2:** Posterior parameter estimates achieved using SCR data alone or integrated with other systematic (telemetry) and opportunistic (opp) information available for the brown bear population in the Italian Alps. Parameters are denoted as follows: number of males and females, *N*_*m*_ and *N* _*f*_, respectively; scale parameter of the Gaussian kernel, which relates to space usage of females and males, *σ* _*f*_ and *σ*_*m*_, respectively; baseline encounter probability (*p*_0_) intercept, *γ*_0_; effect of being male on *p*_0_, *γ*_1_; effect of trap type on *p*_0_, *γ*_2_; effect of time since last check on *p*_0_, *γ*_3_; density intercept, *β*_0_; effect of distance from the release point on density, *β*_1_; data augmentation parameter, *ψ*; probability of being a male, *ω_sex_*. Estimates are reported for a 4-km resolution of the state space grid.

| Parameter | mean | SD | 2.5% | 50% | 97.5% |
| --- | --- | --- | --- | --- | --- |
| SCR + tel + opp |  |  |  |  |  |
| $N_m$ | 13.088 | 3.893 | 6.840 | 12.629 | 21.953 |
| $N_f$ | 26.587 | 10.249 | 13.275 | 24.578 | 51.722 |
| $\sigma_m$ | 10.372 | 0.474 | 9.496 | 10.351 | 11.348 |
| $\sigma_f$ | 8.548 | 0.481 | 7.669 | 8.523 | 9.558 |
| $\gamma_0$ | -3.714 | 0.308 | -4.335 | -3.705 | -3.125 |
| $\gamma_1$ | 1.623 | 0.312 | 1.024 | 1.618 | 2.246 |
| $\gamma_2$ | -1.153 | 0.387 | -1.899 | -1.155 | -0.384 |
| $\gamma_3$ | 0.619 | 0.185 | 0.248 | 0.621 | 0.976 |
| $\beta_0$ | -3.245 | 0.670 | -4.719 | -3.192 | -2.081 |
| $\beta_1$ | -1.128 | 0.467 | -2.125 | -1.107 | -0.276 |
| $\psi$ | 0.198 | 0.064 | 0.109 | 0.186 | 0.351 |
| $\omega_{sex}$ | 0.341 | 0.096 | 0.171 | 0.335 | 0.542 |
| SCR + opp |  |  |  |  |  |
| $N_m$ | 13.687 | 4.297 | 6.965 | 13.166 | 23.675 |
| $N_f$ | 36.890 | 18.900 | 15.703 | 32.157 | 89.949 |
| $\sigma_m$ | 11.800 | 0.905 | 10.207 | 11.741 | 13.743 |
| $\sigma_f$ | 7.092 | 1.103 | 5.380 | 6.936 | 9.669 |
| $\gamma_0$ | -3.414 | 0.345 | -4.116 | -3.406 | -2.758 |
| $\gamma_1$ | 1.417 | 0.360 | 0.734 | 1.409 | 2.143 |
| $\gamma_2$ | -1.160 | 0.392 | -1.923 | -1.164 | -0.390 |
| $\gamma_3$ | 0.621 | 0.189 | 0.248 | 0.623 | 0.985 |
| $\beta_0$ | -2.713 | 0.600 | -3.960 | -2.687 | -1.562 |
| $\beta_1$ | -0.739 | 0.435 | -1.620 | -0.731 | 0.113 |
| $\psi$ | 0.252 | 0.106 | 0.124 | 0.227 | 0.542 |
| $\omega_{sex}$ | 0.291 | 0.096 | 0.125 | 0.285 | 0.498 |
| SCR + tel |  |  |  |  |  |
| $N_m$ | 15.514 | 5.297 | 7.747 | 14.684 | 28.297 |
| $N_f$ | 28.206 | 11.284 | 13.501 | 25.819 | 57.590 |
| $\sigma_m$ | 8.556 | 0.425 | 7.773 | 8.538 | 9.441 |
| $\sigma_f$ | 8.339 | 0.482 | 7.461 | 8.316 | 9.345 |
| $\gamma_0$ | -3.479 | 0.332 | -4.139 | -3.478 | -2.833 |
| $\gamma_1$ | 1.791 | 0.343 | 1.119 | 1.787 | 2.472 |
| $\gamma_2$ | -1.207 | 0.402 | -1.991 | -1.207 | -0.413 |
| $\gamma_3$ | 0.670 | 0.193 | 0.288 | 0.671 | 1.046 |
| $\beta_0$ | -2.956 | 0.627 | -4.313 | -2.899 | -1.869 |
| $\beta_1$ | -0.890 | 0.460 | -1.842 | -0.866 | -0.043 |
| $\psi$ | 0.218 | 0.074 | 0.115 | 0.204 | 0.406 |
| $\omega_{sex}$ | 0.364 | 0.098 | 0.187 | 0.359 | 0.569 |
| SCR |  |  |  |  |  |
| $N_m$ | 16.574 | 5.766 | 7.994 | 15.679 | 30.349 |
| $N_f$ | 65.057 | 31.086 | 24.285 | 58.008 | 147.788 |
| $\sigma_f$ | 3.355 | 0.582 | 2.460 | 3.274 | 4.729 |
| $\sigma_m$ | 9.796 | 1.277 | 7.739 | 9.644 | 12.756 |
| $\gamma_0$ | -2.220 | 0.407 | -3.040 | -2.213 | -1.451 |
| $\gamma_1$ | 0.804 | 0.441 | -0.046 | 0.799 | 1.683 |
| $\gamma_2$ | -1.207 | 0.421 | -2.037 | -1.204 | -0.389 |
| $\gamma_3$ | 0.658 | 0.203 | 0.263 | 0.656 | 1.060 |
| $\beta_0$ | -2.076 | 0.549 | -3.259 | -2.032 | -1.119 |
| $\beta_1$ | -0.459 | 0.387 | -1.267 | -0.437 | 0.244 |
| $\psi$ | 0.406 | 0.168 | 0.177 | 0.371 | 0.850 |
| $\omega_{sex}$ | 0.223 | 0.084 | 0.090 | 0.212 | 0.414 |

**Table 3:** Gain in precision of parameter estimates expressed as per cent reduction in the SD of the posterior estimates achieved by integrating different data types, compared to those obtained using classical SCR models. Population size (*N*_*m*_ and *N* _*f*_) and the scale parameter of the observation model (*σ*_*m*_ and *σ* _*f*_) are reported for males and females respectively.

| | $N_m$ | $N_f$ | $\sigma_m$ | $\sigma_f$ |
| --- | --- | --- | --- | --- |
| SCR + telemetry | 8% | 63% | 67% | 17% |
| SCR + opportunistic | 26% | 39% | 29% | -90% |
| SCR + telemetry + opportunistic | 33% | 67% | 63% | 18% |

**Figure 4:**
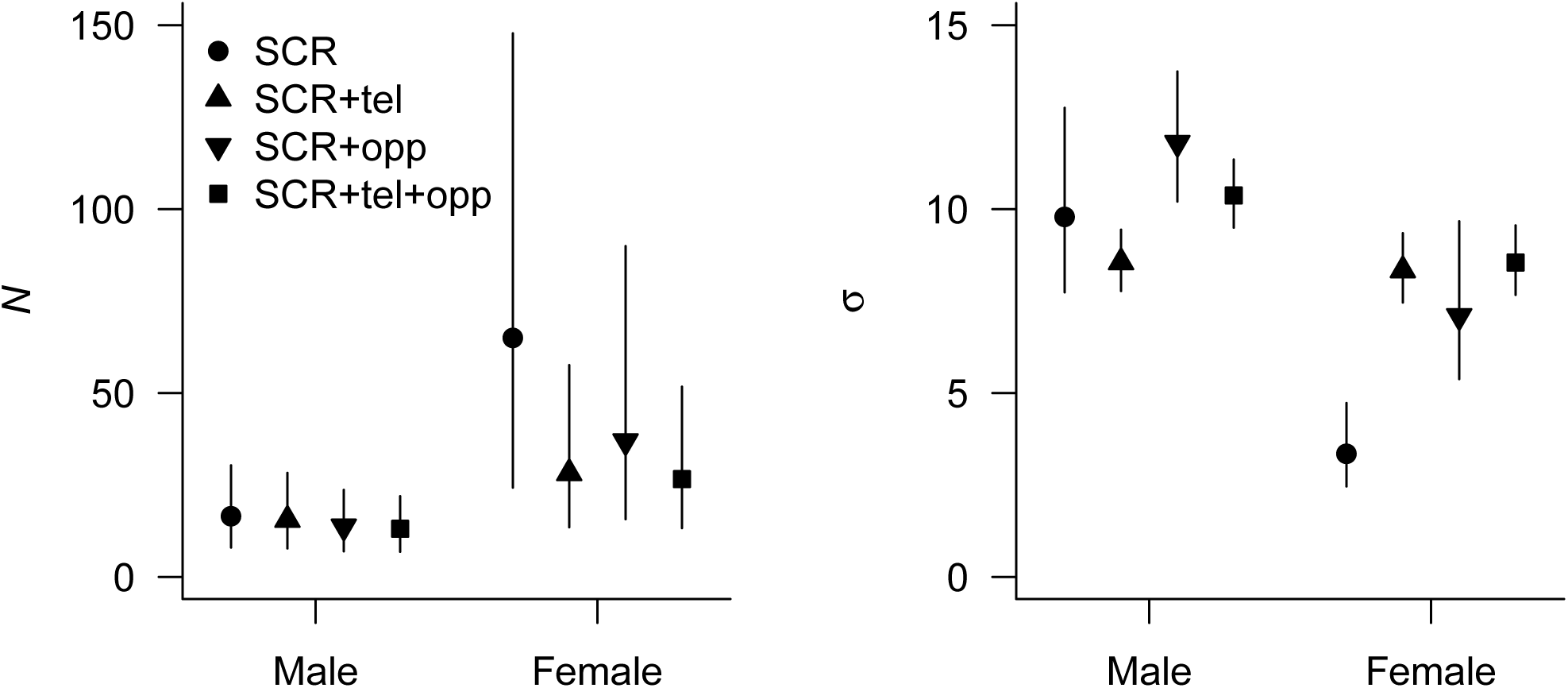
Posterior parameter estimates (mean and 95% Bayesian Credible Interval) achieved using SCR data only, or integrating them with telemetry (‘tel’) and opportunistic (‘opp’) data available for the brown bear population in the Italian Alps. Reported are the sex-specific population sizes (*N*) and scale parameters of the Gaussian kernel (*σ*).

Estimates of the parameter relating density to distance to the reintroduction point (*β*_1_) were negative under all models, and although there was some variation in the strength of the effect, this result supports the hypothesis that density decreased with distance from the point were founders were released (Table 2). The estimated sex ratio in the population, *ω*_*sex*_ did not vary significantly between the four model based on 95% Bayesian Credible Intervals and ranged from 0.22 (BCI: 0.09 – 0.41) in the fully integrated model, to 0.36 (BCI: 0.19 – 0.57) in the SCR + telemetry model (Table 2). Across all models, detectability was higher for males, higher at hair traps, and increased with increasing time between checks (Table 2).

## DISCUSSION

We developed a formulation of a spatial capture-recapture model that integrates multiple data sources, and as a result, improves inferences about key ecological parameters, namely density and space use, which we demonstrate using data from a reintroduced population of brown bears in the Italian Alps. Specifically, we were able to jointly analyze traditional SCR data resulting from systematic sampling of hair traps and rub trees, satellite telemetry data, and opportunistic recovery of biological samples. Comparing estimates from models ranging from traditional SCR, to a fully integrated SCR model that includes both telemetry data and opportunistic sightings data, we demonstrated that the addition of unstructured data results in increased precision in estimates of population size and space usage for this species of conservation interest.

Estimates of male population size were stable across all models and precision was highest for the fully integrated model, i.e., the model with most spatial data, and lowest for the SCR-only model, which had the fewest spatial locations (Fig. 4). For females, estimates of population size from models with additional data were comparable but were different from the SCR-only model estimates, both in terms of precision and point estimates. The addition of telemetry and opportunistic data reduced parameter uncertainty when compared to the SCR-only model but overall precision was lower for the females. Although the number of individuals observed for the two sexes was similar (12 females and 10 males), the number of SCR encounters at different sites (hair traps and rub trees) was smaller for females (mean 2.1, min 1, max 4) than males (mean 8.6, min 1, max 21) (Fig. S1 in Appendix S1). This suggests that smaller SCR data sets might benefit most from an integrated data approach, or conversely may be less stable and more sensitive to combining data. It is encouraging however, that population size estimates based on the two independent unstructured data sources produce similar estimates of abundance, suggesting the addition is beneficial rather than due to parameter sensitivity. The ability to model sex-specific effects on detectability, which in turn affect estimates of sex-specific abundance, can often be limited by insufficient observations of one sex or the other (Sollmann et al. 2011; Tobler and Powell 2013). In cases like these, including in our study, even small amounts of unstructured data, like opportunistic sightings data, can resolve such limitations and increase the value of small SCR data sets.

As with abundance, estimates of sex-specific *σ* were more precise when more data were used, such that *σ* under the fully integrated model had higher precision compared to the SCR-only model (Fig. 4). The most notable difference was between point estimates of female space use from the SCR-only model and the three comparable data integration models. The change in the point estimates is due to the marked difference in the spatial distribution of telemetry locations compared to the SCR data (hair traps and tree rubs) which is likely related to the link between detector type and behavior. SCR data was represented by a few spatially clustered encounters which may mostly reflect female hair deposition patterns related to territoriality potentially at the core of a home range, telemetry and opportunistic data reflect overall space use (Fig. 2b). Estimates of abundance are explicitly linked to estimates of space use because *σ* controls overall expected encounter probabilities. Here, with the integration of added spatial information, estimates of *σ* are higher compared to SCR-only data, and as a result, estimates of abundance are reduced (Fig. 4). Again, this points to the value of added spatial information to refine estimates of space use, and in turn, of abundance and density.

The integration of additional spatially-referenced information with traditional SCR data ameliorate inference accuracy, and even a small amount of opportunistic data yields improvements, meaning that citizen science type data are potentially high value data sets in SCR regardless of whether survey effort is known. In some cases, like for the male movement parameter *σ_m_*, incorporation of opportunistic information may counterbalance the possible inconsistency between spatial information provided by telemetry and SCR data (Fig. 4). This is particularly relevant in studies of rare or elusive species, where the amount of SCR encounters is usually small but also budget restrictions and the difficulty in collaring animals limit the number of individuals for which telemetry information is available. Opportunistic data provide additional information not limited to the extent of a trap array, like telemetry, but which can also be available for larger number of individuals than those equipped with devices. Unsurprisingly, the addition of telemetry data alone generally results in more precision gains than adding opportunistic data alone because the former is more information rich (Fig. 4). However, the addition relatively few opportunistic locations acted to increase precision relative to inference from the SCR-only model (Fig. 4), suggesting that even in the absence of telemetry data, opportunistic data are important sources of spatial information.

When the number of collared individuals is very low (e.g. 3 individuals in Royle et al. 2013 and Sollmann et al. 2013a, or two in our study) telemetry information may be more or less representative of population space use, with a variable degree of concordance between spatial information provided by telemetry and SCR data. On the other hand, information mismatch can also arise in the presence of sparse SCR data as a consequence of inappropriate trap spacing, variation in group-specific (e.g. age or sex) exposure to traps, or when the trap array is small relative to individual movement, inducing a geographic bias for the most mobile component of the population. Opportunistic information may also not be representative of the entire population, as in the case of records collected at sites where a damage occurred (e.g. depredation on livestock, beehives and crops) where individuals more prone to commit damage can be sampled more times than others. Finally, as suggested above, the encounter data may reflect altogether different behavioral states depending on the particular method used to detect individuals. This suggests that the process of data integration requires more that simply the development of integrated data models, but rather that those models take into consideration the variation in behavior that might be reflected in the independent data sets being used.

We provide evidence that incorporating unstructured opportunistic data to SCR and telemetry information, by conceptually treating opportunistic records as thinned telemetry data, improves inference on abundance and space usage, which are key population-level parameters to inform conservation decisions of elusive and difficult-to-study species. However, care must be taken to assess the potential mismatch in spatial information provided by the different data sets, where telemetry is both the most informative source of space use but also often available only for a few individuals, whose movement may not be representative of population space use. Understanding how animal density changes in space and how the latter is used is crucial when addressing practical issues in population management and conservation (Bischof et al. 2009). For this aim, the use of opportunistic information increases availability of spatially-referenced individual information, that can be suitably modelled along with other data within a unified framework, thus reducing the need for additional invasive methods.

## ACKNOWLEDGEMENTS

This research was partially funded by the Autonomous Province of Trento and the MUSE - Museo delle Scienze. We would like to thank the Autonomous Province of Bolzano, the ISPRA, the personnel of the Servizio Foreste e Fauna of the Autonomous Province of Trento, of the Adamello Brenta Natural Park, and of the Stelvio National Park. We also thank the many forestry wardens and volunteers for field support, and Aaron Iemma for IT assistance.

